# The epigenomic landscape of deep lineage divergence: The case of the European sea bass

**DOI:** 10.64898/2026.07.27.738391

**Authors:** Alessio Longo, Massimiliano Babbucci, Zexin Jiao, Serena Ferraresso, Rafaella Franch, Martina Bortoletti, Daniela Bertotto, Sara Faggion, Garth R. Ilsley, Vasileios Papadogiannis, Tereza Manousaki, Jon Kristoffersen, Costas S. Tsigenopoulos, Daniel J. Macqueen, Luca Bargelloni

## Abstract

**Background:** Understanding the role of non-coding genomic variation in speciation remains a major challenge in evolutionary biology. Here, we investigated whether regulatory elements contribute to this process between Atlantic and Mediterranean lineages of European sea bass (*Dicentrarchus labrax*), a well-characterized case-study near speciation where barriers to introgression exist in the presence of connectivity between diverging populations.

**Results:** We generated a novel, highly contiguous genome assembly, which was annotated at the epigenomic level using ATAC-seq and ChIP-seq with six embryonic developmental stages and five tissue types in adult fish, identifying thousands of promoters, enhancers, and open chromatin regions. Integrating this annotation with whole-genome sequence data from 65 individuals across three geographically distinct populations, we identified 57,505 outlier SNPs and 332 structural variants (SVs) showing elevated differentiation between Atlantic and East Mediterranean lineages. Outlier SVs affected key regulatory elements and coding genes, while outlier SNPs were enriched in regulatory elements, particularly enhancers active in adult tissues. Local genomic divergence correlated positively with regulatory element density, especially on chromosomes 1, 9, and 18, which are enriched in genes related to osmoregulation, immune response, and oxidative stress — processes relevant to adaptation across contrasting marine environments.

**Conclusions:** These findings support a major role for regulatory variation in driving deep lineage divergence through local adaptation.

## Background

It has long been proposed that genetic variants regulating gene expression play an important role in evolution^1^. Since then, a wealth of experimental evidence has supported this hypothesis^2–4^. Non-coding variants primarily shape traits by impacts on gene expression. Unlike coding variants that alter protein function, non-coding variants can modulate how, when, and where genes are expressed, primarily via changes to promoters, enhancers, silencers, and other cis-regulatory elements. Such elements are identifiable in eukaryotic genomes, owing to continuous improvements in genome functional annotation^5–7^. Genome-wide association studies have also demonstrated that most lead variants map to regulatory regions, while expression QTL analyses have revealed impacts on splicing, transcription factor binding, and chromatin structure, all of which are integral to gene regulation and cellular function^8^.

The drivers of speciation have attracted the interest of evolutionary biologists since Darwin. Noncoding variants in regulatory elements play a crucial role, impacting species-specific adaptations and reproductive isolation^9^. The role of gene expression in speciation has been explored by comparing transcriptomic profiles in divergent lineages and their hybrids^10,11^. A complementary strategy focuses on genomic regions of increased differentiation between divergent lineages^12^. The present study adopts this approach to test the hypothesis that non-coding genomic variation and regulatory elements have shaped the divergence of European sea bass (*Dicentrarchus labrax*) populations across the Atlantic-Mediterranean divide.

Several marine species show a clear phylogeographic break between Mediterranean and Atlantic populations^13^. Among these, the European sea bass has been extensively investigated as an exemplar of deep lineage divergence in the marine realm in the absence of strong present-day barriers to gene flow. Divergence between Atlantic and Mediterranean populations originated from allopatric separation accumulated for about 300,000 years^14^. Despite the re-opening of the Gibraltar strait at the end of the last glaciation, and the potential for connectivity between the Atlantic and the Mediterranean, introgression between the two lineages appears limited. A history of isolation during the last glacial maxima, coupled with ancient admixture with a closely related species (*D*. *punctatus*), appears to have generated genomic islands of increased divergence, which were subsequently shaped by further introgression and heterogeneous patterns of recombination^14–16^. Such islands of high genomic divergence may be associated with genomic incompatibilities as well as positive selection^12^. In the case of European sea bass, it was suggested that these islands caused Bateson-Dobzhansky-Muller (BDM) incompatibilities^16^, although a recent study showed that selection against maladapted alleles has shaped the genomic landscape of introgression between Atlantic and Mediterranean lin-eages^17^. In fact, European sea bass geographic populations display heritable phenotypic divergence for different traits such as body shape and growth, thermal growth coefficient, muscle fat content^18^, and disease resistance^19,20^. The introgressed populations in the Western Mediterranean might also have reduced fitness, as they display male-biased sex ratios and lower body weight^21^. Furthermore, as the Atlantic and the Mediterranean basins show significant clines in salinity and water temperature, that adaptation to local osmotic and thermal conditions might contribute to shaping genomic divergence between populations.

In this study, a comprehensive atlas of regulatory elements and gene expression was produced for European sea bass, representing different stages of embryonic development and a panel of adult tissue samples. Specifically, ATAC-seq and ChIP-seq datasets were generated and mapped against a new highly contiguous genome assembly, with all resulting regulatory elements shared through the Ensembl genome browser (https://www.ensembl.org/Dicentrarchus_labrax/Info/Index). Individual whole-genome sequence data^16^ were re-analyzed to identify single nucleotide polymorphisms (SNPs) and structural variants (SVs). Genomic divergence was then assessed in relation to this epigenomic landscape to test whether non-coding variants, in particular those affecting regulatory elements, are involved in deep divergence between the Atlantic and the Mediterranean lineages.

## Results

### The European sea bass genome assembly

The long-read assembly produced was composed of 574 contigs, with an N50 of 12,698,854 bp, a BUSCO score of 95.4%, and 105 missing BUSCOs. Polishing with the short-read data provided an improved assembly with a total length of 694,870,088 bp, an N50 of 12,677,075 bp, increased the BUSCO score to 97.8% and reduced the missing BUSCOs to 63. Further scaffolding of the contigs of this polished assembly, guided by the published assembly by Tine et *al*^15^, provided a final assembly composed of 302 scaffolds, with a total length of 695,892,153 bp, an N50 of 29,885,359 bp. As the 24 largest scaffolds contain 96.669% of the total length of the assembly, this is a near chromosome level assembly, with the 24 largest scaffolds likely corresponding to the 24 chromosomes of the species.

### The European sea bass epigenome landscape

ATAC-seq and ChIP-seq data (for histone marks: H3K4me1, H3K4me3 and H3K27a) were obtained for six embryonic stages (early gastrula, mid gastrula, early somitogenesis, mid somitogenesis, late somitogenesis, near hatching) and six adult tissues (brain, skeletal muscle, gills, ovary, testis, liver). With the obvious exception of gonads, for each adult tissue ATAC-seq, ChIP-seq, and RNA-seq data were obtained independently for four conditions (immature adult males, mature adult males, immature adult females, mature females). For the sake of clarity, overall tissue data are reported here, although condition-specific evidence is available in Ensembl. Integrating this dataset, three types of regulatory elements were identified using the Ensembl regulatory build pipeline described in Ilsley et *al*^22^. A broad gene expression atlas obtained for the same samples using RNA-seq complemented this epigenome profile, providing functional annotation of the sea bass genome spanning both genes and regulatory elements (https://www.ensembl.org/Dicentrarchus_labrax/Info/Index). This epigenome profile was then analyzed against a novel sea bass genome assembly (https://www.ensembl.org/Dicentrarchus_labrax/Info/Annotation; details in Methods).

The minimum number of overall regulatory elements detected was 31,711 at the early gastrula stage (**Table 1**). An increasing trend was observed across development, reaching a maximum during late somitogenesis (**Fig. 1A**) with 53,847 regulatory elements identified (**Table 1**). Enhancers were the most abundant regulatory regions across developmental stages and exhibited a generally linear increase from early gastrula (13,299) to late somitogenesis (25,515), followed by a slight decrease at the final developmental stage (22,174). A similar pattern was observed for open chromatin elements, which increased from 12,020 at the early gastrula stage to 21,065 during late somitogenesis stage, before slightly decreasing to 18,044 as eggs approached hatching. In contrast, promoters showed a less linear trend, with a slight decrease from early to mid-gastrula, an increase at early somitogenesis followed by a decrease at mid-somitogenesis, an increase during late somitogenesis, and a final decrease at the last developmental stage. Stage-specific elements ranged from 507 at mid-somitogenesis to 2,984 at the early gastrula stage (**Table 1**) and displayed a distinct pattern, with the highest number of regulatory elements detected during early gastrulation (**Table 1**, **Fig. 1B**). Overall, open chromatin elements were the most frequently identified across stages, followed by enhancers and promoters.

**Fig. 1.**
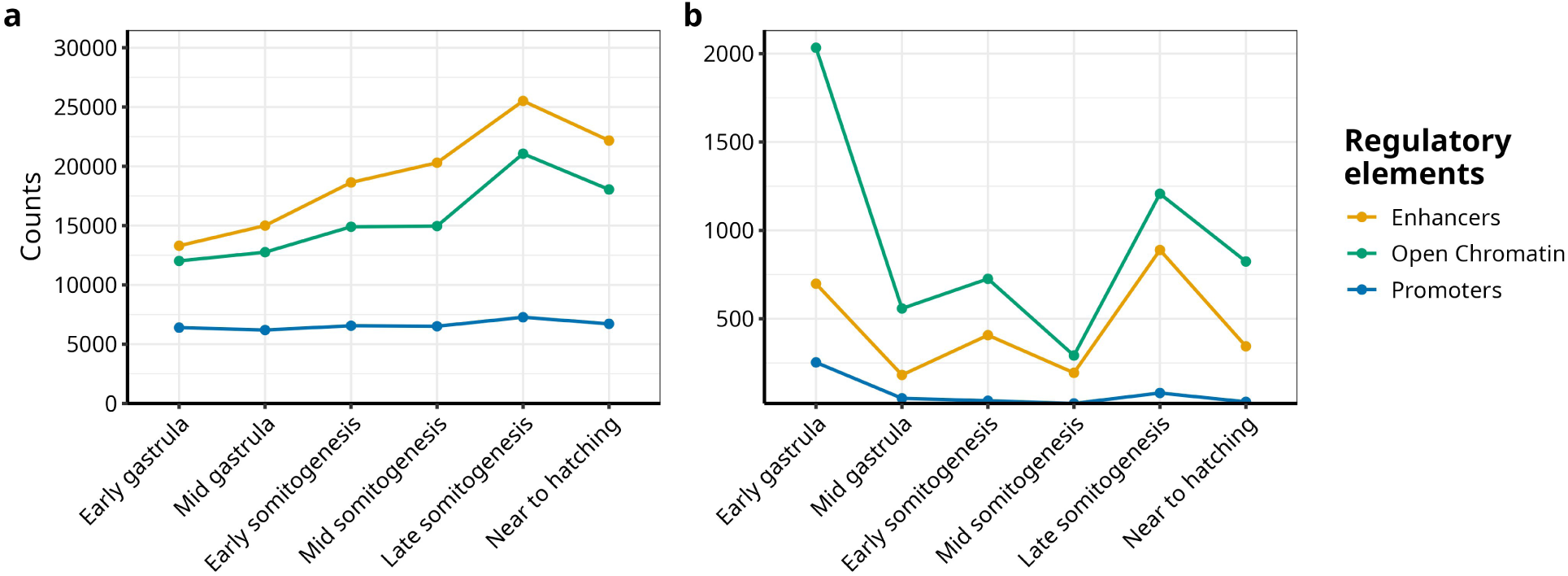
Patterns of active regulatory element during early development. Trends in active regulatory elements observed across developmental stages showing **A**, overall count and **B**, stage-specific regulatory elements. Enhancers, open chromatin regions, and promoters are depicted by yellow, green and blue lines, respectively.

**Table 1.** Number of overall and stage-specific regulatory elements per type and developmental stage.

| Develop-<br>mental<br>stage | Overall<br>regula-<br>tory el-<br>ements | Pro-<br>moters | En-<br>hance<br>rs | Open<br>Chro-<br>matin | Stage-spe-<br>cific regu-<br>latory el-<br>ements | Pro-<br>moters | En-<br>hancers | Open<br>Chro-<br>matin |
| --- | --- | --- | --- | --- | --- | --- | --- | --- |
| Early gas-<br>trula | 31,711 | 6,392 | 13,299 | 12,020 | 2,984 | 253 | 698 | 2,033 |
| Mid Gas-<br>trula | 33,949 | 6,192 | 15,000 | 12,757 | 790 | 50 | 182 | 558 |
| Early somi-<br>togenesis | 40,093 | 6,553 | 18,637 | 14,903 | 1,169 | 36 | 407 | 726 |
| Mid somi-<br>togenesis | 41,771 | 6,509 | 20,302 | 14,960 | 507 | 21 | 194 | 292 |
| Late somi-<br>togenesis | 53,847 | 7,267 | 25,515 | 21,065 | 2,176 | 80 | 889 | 1,207 |
| Near to<br>hatching | 46,933 | 6,715 | 22,174 | 18,044 | 1,198 | 30 | 344 | 824 |

In adult tissues, the number of all categories of active regulatory elements was perfectly correlated with the level of tissue-specificity (Spearman ρ = 1, p<0.0000001), with the lowest number of elements active in all six tissues and the highest number for elements active in a single tissue (**Table 2**).

**Table 2.** Number of overall regulatory elements in tissues collected from immature and mature adult males and females.

| Tissue-specificity | Overall regulatory elements | Promoters | Enhancers | Open Chromatin |
| --- | --- | --- | --- | --- |
| Six tissues | 3,922 | 1,403 | 2,382 | 137 |
| Five tissues | 8,287 | 2,817 | 4,828 | 642 |
| Four tissues | 11,702 | 3,586 | 6,720 | 1,396 |
| Three tissues | 17,228 | 4,623 | 9,854 | 2,751 |
| Two tissues | 29,122 | 5,954 | 17,260 | 5,908 |
| One tissue | 81,349 | 9,372 | 47,608 | 24,369 |

The tissues with the highest number of strictly tissue-specific elements were gills, liver, and brain (13,433, 10,609 and 8,658, respectively; **Table 3**). Within these tissues, enhancers were the most frequently identified regulatory elements, ranging from 6,221 in brain to 7,094 in liver, followed by open chromatin elements (from 1,906 in brain to 6,247 in gills). Promoters were the least abundant elements, ranging from 484 in gills to 851 in liver. In contrast, female and male gonads, as well as skeletal muscle, were characterized by lower numbers of regulatory elements (2,548, 1,064 and 1,248, respectively). Similar to brain, gills and liver, female gonads and muscle exhibited a predominance of enhancers and relatively few promoters. In male gonads, open chromatin elements were the most abundant category, followed by enhancers and promoters.

**Table 3.**
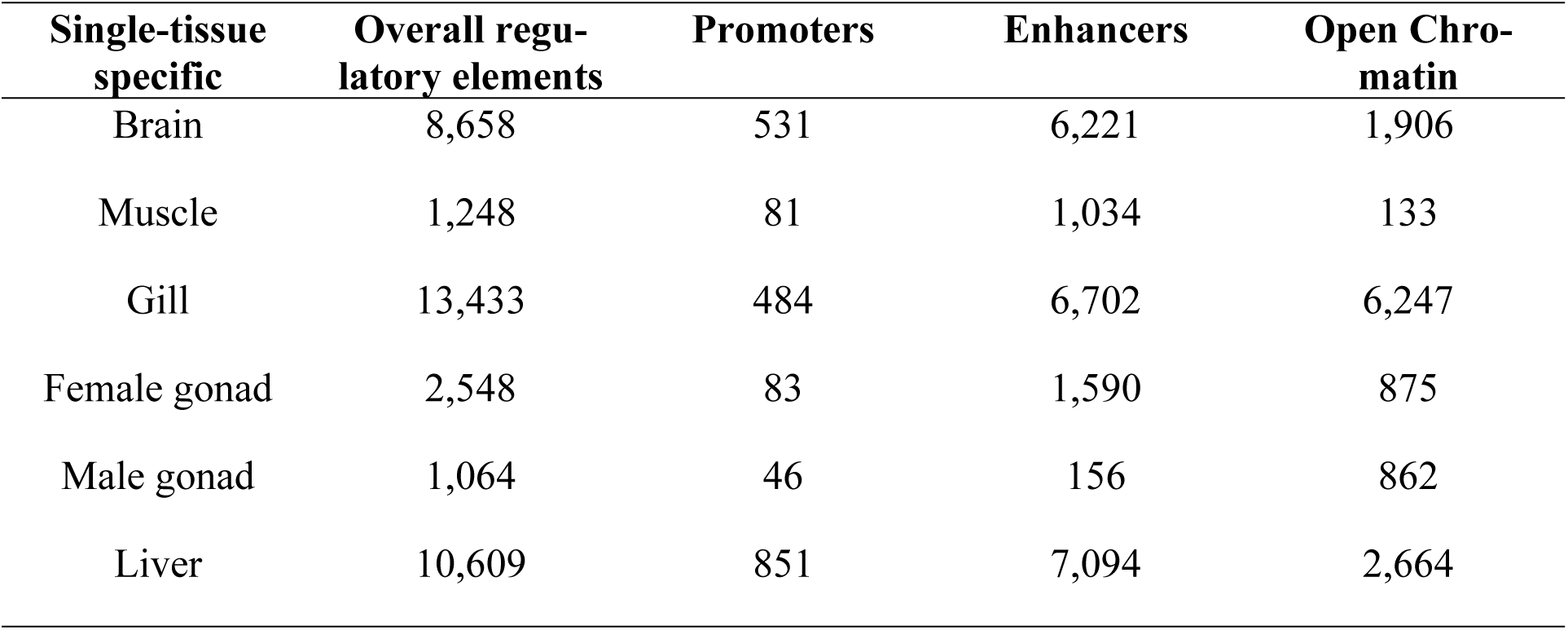
Number of tissue-specific regulatory elements for each category of tissue-specificity.

| Single-tissue specific | Overall regulatory elements | Promoters | Enhancers | Open Chromatin |
| --- | --- | --- | --- | --- |
| Brain | 8,658 | 531 | 6,221 | 1,906 |
| Muscle | 1,248 | 81 | 1,034 | 133 |
| Gill | 13,433 | 484 | 6,702 | 6,247 |
| Female gonad | 2,548 | 83 | 1,590 | 875 |
| Male gonad | 1,064 | 46 | 156 | 862 |
| Liver | 10,609 | 851 | 7,094 | 2,664 |

The distribution of elements along the genome appears rather uniform, as the mean percentage of genome annotated as part of a regulatory element is similar across all 24 sea bass chromosomes, with only two outliers for open chromatin regions (Chromosome 15 and 22, **Additional file 1: Table S1**) identified by Tukey’s fences test. The distribution of mean coverage per element type is shown in **Fig. 2**.

**Fig. 2.**
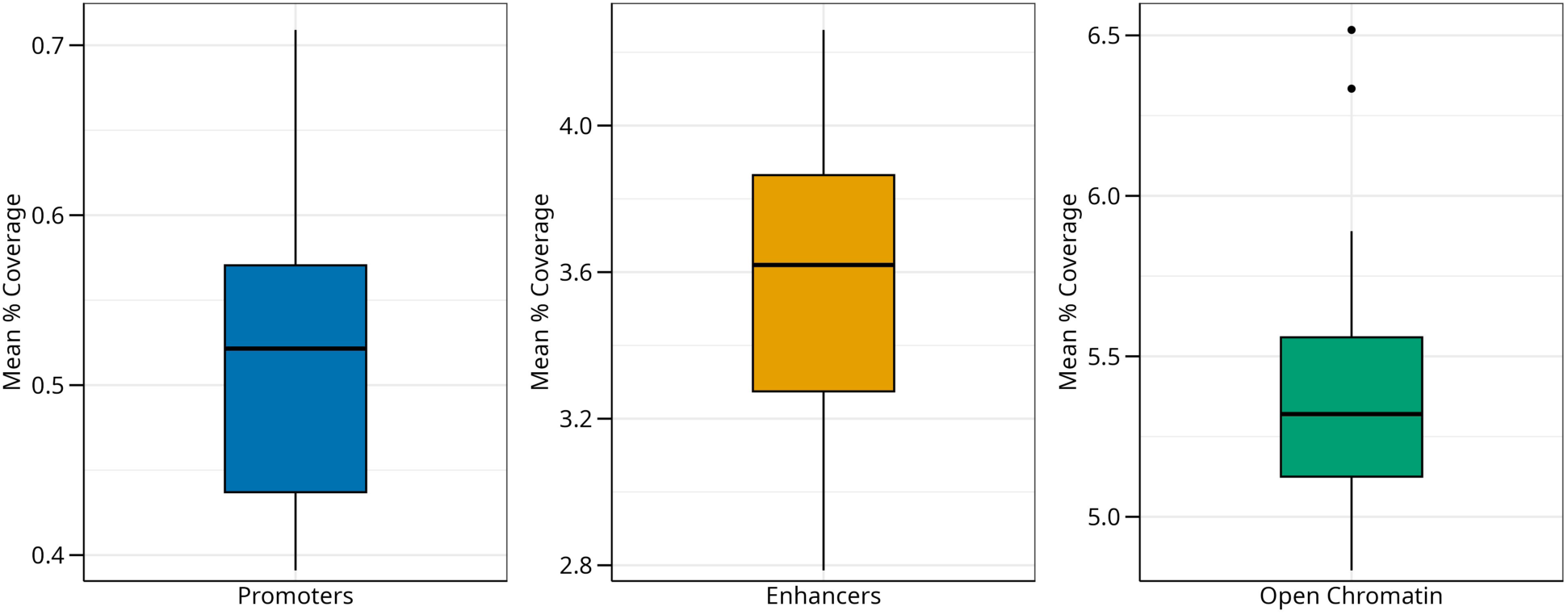
Boxplots of mean percentage of each chromosome annotated as promoters, enhancers, and open chromatin. Blue, yellow and green boxplots represent mean coverage percentages of promoters, enhancers and open chromatin, respectively.

### Genomic divergence between Atlantic and Mediterranean lineages at non coding sites

Whole genome sequence data for 65 unrelated individuals from three different geographic areas (Atlantic, West Mediterranean, andEast Mediterranean) were considered. Only the 24 largest scaffolds, corresponding to the 24 sea bass chromosomes were taken forward in analyses. After filtering for minor allele frequency, a total of 7,128,064 SNPs and 38,408 SVs were retained. The SVs were predominantly deletions (37,074, >96%), while 1024 duplications and 310 inversions were detected. Cluster analysis using fastSTRUCTURE v1.0^23^ was performed to identify individuals showing minimal admixture between the Atlantic and Mediterranean lineages (**Additional file 2: Fig. S1**), confirming previous evidence that Atlantic and East Mediterranean samples are highly divergent, while West Mediterranean individuals show variable levels of introgression. Genomic variation between the most divergent groups was analyzed using pcadapt to reveal outlier loci showing significantly higher divergence. After Bonferroni correction, 57,505 significantly divergent SNPs and 332 SVs were recovered. A Fisher’s exact test indicated that SNPs were significantly overrepresented in the outlier category, while SVs were underrepresented (p<1*10^-12^).

Outlier SNPs were non-randomly distributed across the genome. SNPs located in regulatory elements were compared with SNPs present in genomic regions that are at least 3kb either upstream or downstream annotated genes and do not overlap regulatory elements. We assumed that SNPs located in these regions are less functionally relevant. Such comparison showed that outlier SNPs were significantly enriched in regulatory elements (χ2 p<1*10^-5^). Among the three types of elements, SNPs in enhancers were significantly overrepresented (χ2 p<1*10^-5^). Outlier SNPs were also overrepresented in gene bodies compared to intergenic regions (χ2 p<1*10^-4^). Within gene bodies, comparing synonymous and non-synonymous SNPs revealed no significant enrichment of either category in outlier loci, while the comparison of intronic SNPs (excluding intronic regulatory elements) and SNPs precisely located in splice sites showed marginally significant enrichment of outlier SNPs affecting splice sites (χ2 p<0.05). Likewise, SNPs located within gene bodies either in 5’ or 3’ UTRs or in coding regions were significantly enriched in outliers compared to introns (χ2 p<0.01), after excluding regulatory elements in all these regions. Functional analysis of genes harboring outlier SNPs provided only marginally significant enrichment for broad GO categories (data not shown). Specific enrichment analysis for nuclear genes encoding mitochondrial proteins (MitoCarta3.0) showed no significant results (**Additional file 1: Table S2**).

### Outlier SVs analysis

We identified 15 outlier deletions affecting mature transcripts (5’ or 3’ UTRs or coding regions), significantly more than expected compared to those within introns (χ2 p<1*10-4). Of those 15 deletions, five are predicted to create truncated forms of the encoded protein. A relatively large deletion (>700 bp) ablated either one or two downstream coding exons, depending on isoform, of gene EN-SDLAG00005004700, encoding the orthologue of human solute carrier family 22 member 4 (SLC22A4) (**Fig. 3A**).

**Fig. 3.**
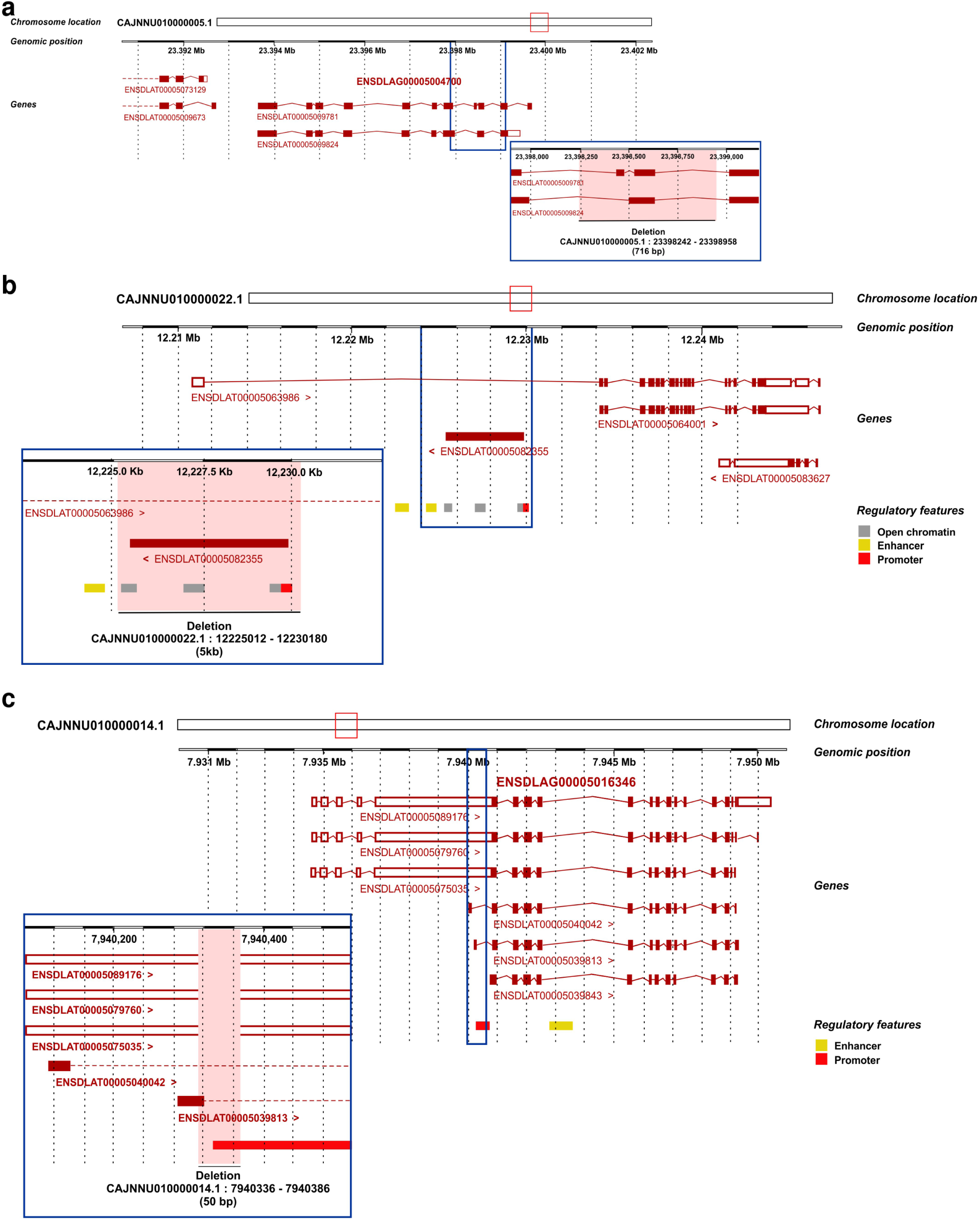
Notable examples of outlier SVs. **A** Large deletion SV ablating coding exons of ENSD-LAG00005004700. **B** Genomic structure of the >5kb deletion on chromosome 22, removing four open chromatin regions and one promoter. **C** A 50 bp deletion SV disrupting the promoter element ENSDLAR00000189036, involved in the alternative transcript ENSDLAT00005039813.2 of the guanidine deaminase gene ENSDLAG00005016346

A smaller deletion (<100 bp) is predicted to encode the second exon of gene EN-SDLAG00005031078, encoding the putative orthologue of flavin-associated monooxygenase 5, while another small SV shortened the first exon of the uncharacterized gene EN-SDLAG00005025579. A 50 bp deletion truncated the first exon of one of the alternative transcripts (ENSDLAT00005039813.2) of gene ENSDLAG00005016346, encoding guanidine deaminase. Finally, a larger SV deleted the first coding exon of gene ENSDLAG00005004017, which encodes cadherin 26.

A total of 23 outlier SVs, 19 deletions and 4 duplications, overlapped annotated regulatory regions. These SVs affected 19 open chromatin regions, 8 enhancers, and 2 promoters. The number of tissues, conditions, or stages where the affected regulatory elements are active varied from 1 to 23, with a median value of 3. A few deletions and duplications affected more than one element. The most remarkable example is a large deletion (>5kb) located on chromosome 22, which removes three open chromatin regions and one promoter (ENSDLAR00000218305, ENSDLAR00000218306, EN-SDLAR00000218307, ENSDLAR00000218308) (**Fig. 3B**). All four elements are active in the mature male gonad and overlap with an LTR-retrotransposon of the Gypsy11 subfamily. It is therefore likely that this large deletion is due to the presence or absence of the entire gypsy-like elements. The other SV affecting a promoter has been already described above. This element (EN-SDLAR00000189036) appears to be a liver-specific alternative promoter immediately upstream of the shortest isoform of gene ENSDLAG00005016346, encoding guanidine deaminase. The promoter element located upstream to all other alternative transcripts (ENSDLAR00000189033) is active in a much broader set of tissue-stages (brain, all embryonic stages, skeletal muscle, gills, male and female gonads, liver) (**Fig. 3C**).

### Outlier SNP analysis

Enhancers appear to harbour significantly more outlier SNPs than expected compared to intergenic regions. Therefore, we tested the effect of either the stage(s) or the tissue(s) in which these regulatory elements are active. Outlier SNPs were observed more frequently in enhancers active only in adult tissues, compared to those active either only in embryonic stages or both in embryos and adult stages (χ2 p<1*10^-9^). No single embryonic stage or adult tissue showed significant outlier SNP enrichment in stage-or tissue-specific enhancers. Similarly, breath of activity across adult tissues (from single-tissue specific enhancers to regulatory elements active in all tissues) showed no association with outlier SNP abundance.

A complementary analysis to test the association between functional features and genomic divergence was carried out by correlating median Fst values and the coverage of either gene bodies or different types of regulatory elements in non-overlapping 100kb windows along the 24 chromosomes. Overall, the percentage of gene bodies per window showed a weak positive correlation with corresponding Fst values in the same window (Spearman ρ = 0.05, p<0.00001). Higher and more significant positive correlations were observed for the percentage of promoter regions (Spearman ρ = 0.13, p<1*10^-15^) and enhancers (Spearman ρ = 0.125, p<1*10^-14^). The same test on individual chromosomes revealed several cases of highly significant positive correlations (**Additional file 1: Table S3**). For chromosomes 1, 9, and 18, the presence of both promoters and enhancers was positively correlated with Fst (ρ with range 0.21-0.32) (**Fig. 4**).

**Fig. 4.**
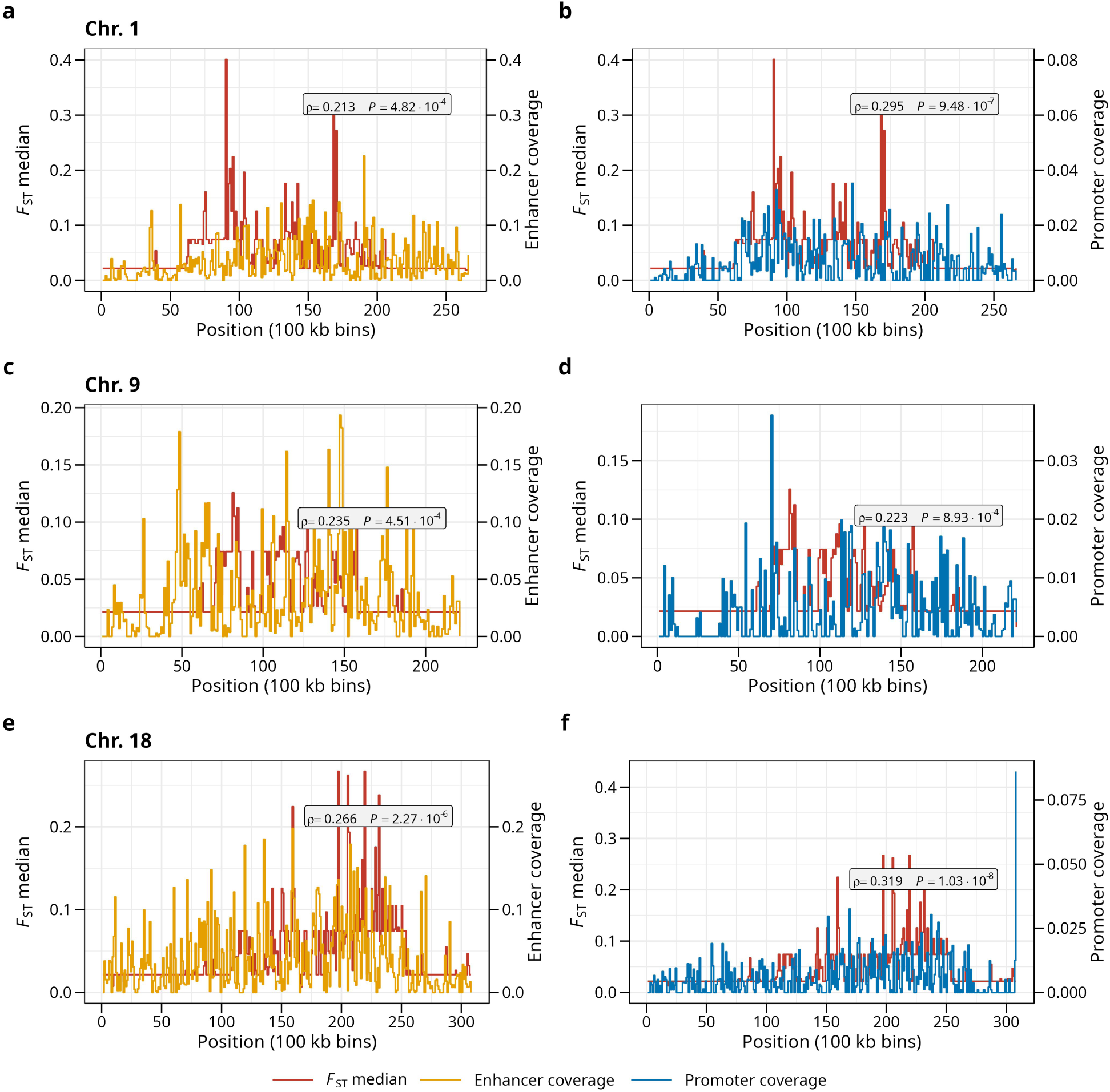
Fst and percentage of genomic regions occupied by enhancers and promoters in 100kb windows along chromosomes 1, 9, and 18. Step charts overlapping median Fst values (red lines) with percentage of enhancer (yellow lines) and promoter coverage (blue lines) in 100 kb bins for chromosome 1 (**A**, **B**), chromosome 9 (**C**, **D**), and chromosome 18 (**E**, **F**), respectively. Grey labels display corresponding Spearman ρ and p-values.

SNPs density was not correlated with the presence of functional features with the only exception of chromosome 17, which showed a significantly positive correlation for both gene bodies and regulatory elements (**Additional file 1: Table S4**), while SVs were overall negatively correlated with the density of either gene bodies or regulatory elements (**Additional file 1: Table S5**).

Functional analysis with g:Profiler showed that chromosomes 1, 9 and 18 are significantly enriched in genes encoding proteins involved in symporter activity exchanging amino acids and either sodium or chloride (chromosome 1, **Fig. 5A**), in positive regulation of immune response (chromosome 9, **Fig. 5B**), and in flavin-associated monooxygenase activity (chromosome 18, **Fig. 5C**), respectively.

**Fig. 5.**
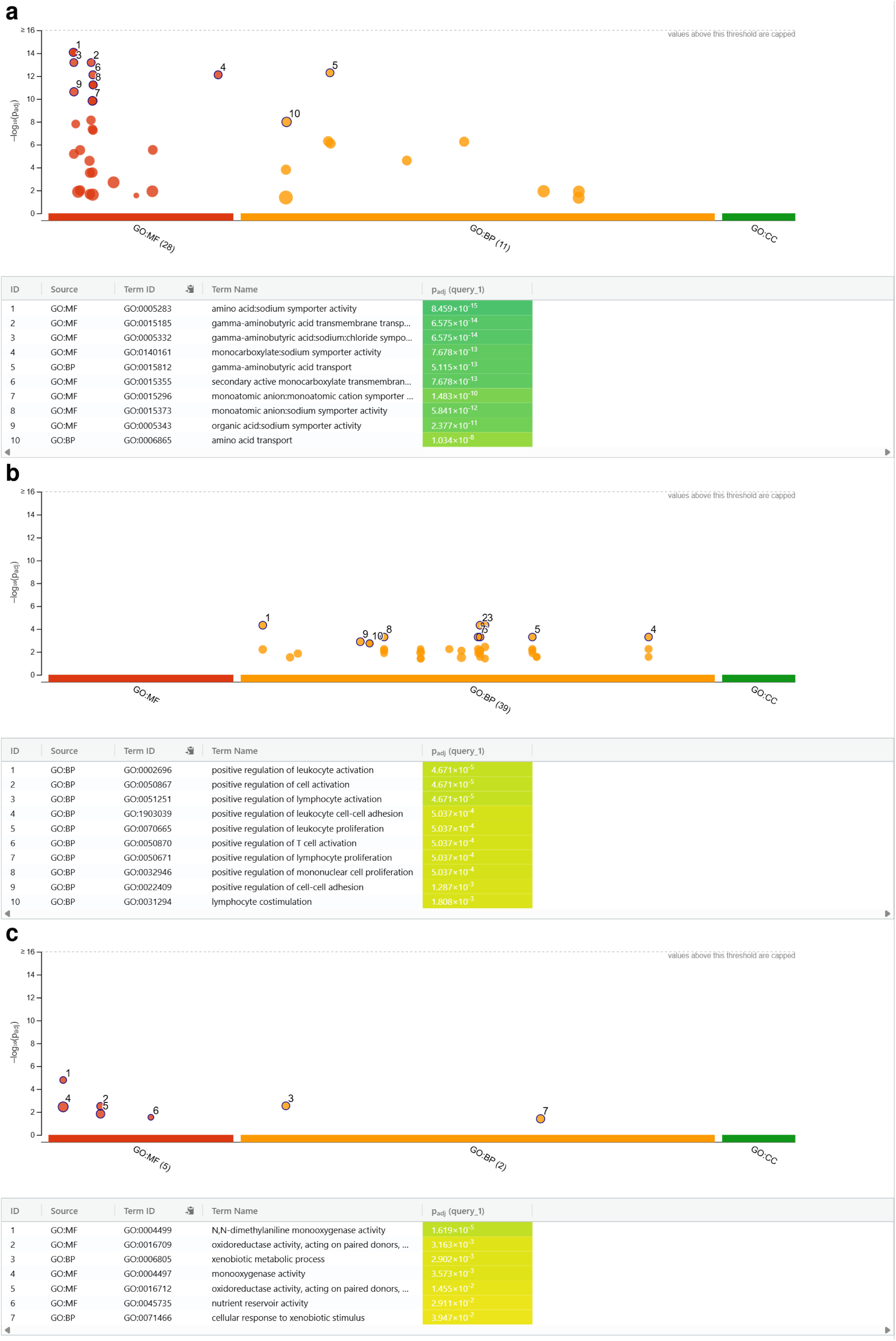
Functional enrichment analysis of chromosomes 1, 9, and 18. Manhattan plots illustrating the enrichment analysis results for **A** chromosome 1, **B** chromosome 9, and **C** chromosome 18. The x-axis shows functional gene ontology (GO) categories clustered and coloured in molecular function (MF; red), biological process (BP; orange), and cellular component (CC; green). The y-axis illustrates the adjusted enrichment p-values in negative log_10_ scale. Below each graph a summary table is presented, detailing the identified specific GO term ID with the associated p-value. For chromosomes 1 and 9 the first ten GO terms are highlighted, while for chromosome 18 all seven identified GO terms are presented.

### Local recombination rate and functional features

Heterogeneity in recombination rates across the sea bass genome may significantly affect analysis of outlier loci and their association with functional genomic features. Local recombination rates were estimated based on a novel linkage map built using SNP-array genotype data (see Methods) from a multifamily panel. Using non-overlapping 100kb windows, median Fst values and local recombination rate per window were negatively correlated across the genome (Spearman ρ =-0.26, p<1*10^-14^). Correlation analysis by chromosome confirmed such evidence (**Additional file 1: Table S6**), while coverage by functional features (gene bodies, promoters, enhancers, open chromatin) appeared to be either only weakly or not correlated with local recombination rate. Just a few chromosomes displayed mostly positive ρ values (**Additional file 1: Table S7**). Density of both types of variants, SNPs and SVs, was positively correlated with local recombination rates (ρ = 0.61844, p<1*10^-14^ and ρ = 0.15, p<1*10^-14^; **Additional file 1: Table S2**).

## Discussion

The main goal of the present paper was to obtain accurate functional annotations of the European sea bass genome as a framework to interpret genomic divergence linked to the Atlantic-Mediterranean divide. A large number of regulatory elements were identified across developmental stages and adult tissues. Adult tissues with the highest number of active elements, gills, liver and brain, are known to have a complex set of cell types, expected to produce a larger number of regulatory elements in bulk epigenomic profiles. While a rigorous comparison of the sea bass epigenome against other verte-brates is beyond the scope of the present study, the number of regulatory elements and their sample-specific activity profiles appear in agreement with other species. For instance, in an in-depth functional annotation of zebrafish *Danio rerio*, Baranasic et *al*^24^ revealed a similar increasing trend in active enhancer elements from the gastrula stage (*75%-epiboly*) to early somitogenesis (*5-9 somites*) peaking at late somitogenesis (*prim-5*), followed by a decrease near hatching (*long-pec*), as also observed in sea bass embryos (**Fig. 1A**).

With regard to the genomic divergence between Atlantic and Mediterranean sea bass lineages, the first important evidence is the bias toward SNPs in the outlier group compared to SVs. It is possible that methodological limitations due to the use of short read sequencing data might have reduced the ability of identifying complex SVs. Comparisons of SVs generated by reference-based short-read and long-read WGS on the same individuals have consistently observed that long-read WGS captures more than double the total number of SVs in the human genome, but the vast majority (>91%) of these novel variants uniquely detected by long-read WGS are located to non-coding repetitive sequences. Conversely, long-read and short-read WGS exhibit strong concordance (>93%) for deletions in the remaining 90% of the genome comprised of sequences that are less repetitive^25^. In the sea bass, the vast majority of SVs are deletions behaving as biallelic, which, together with the relatively large number of identified SVs, should exclude that outlier detection methods optimized for large data sets of biallelic loci might be affected by systematic biases when analyzing SVs. A more likely hypothesis is that selective forces acting on SVs and SNPs of genomic variants are substantially different.

Structural variants, often affecting larger genomic regions, may be less likely to be positively selected because they can generally have significant deleterious effects. Population genetic analysis of SVs in humans suggest that deletions are mostly under purifying selection^25^. Conversely, SNPs might be significantly involved in fine-scale adaptive evolutionary processes, especially in traits under local selection where small-effect mutations accumulate rapidly. On the other hand, deletions involving either entire or large portions of regulatory regions in most cases should significantly affect these elements, likely causing function loss. Likewise, duplications might profoundly alter gene expression regulation^26^. Therefore, most SVs involving regulatory regions are likely under strong purifying selection. However, over 300 SVs were identified as outliers with alternative alleles being nearly fixed respectively in Atlantic and Mediterranean samples. It is difficult to conclude that these SVs behave as either neutral or mildly deleterious mutations. Many of them potentially affect transcriptional regulation, with significant phenotypic effects. Deletions causing truncated protein isoforms might have a strong phenotypic effect. Consequently, while the number of involved loci is much smaller than in the case of SNPs, SVs might well have an important role in the divergence between European sea bass lineages. Using the same approach implemented here, it has recently been proposed that SVs had a role in domestication of European sea bass, as a significant “brain” signature was shown to characterize SVs differentiating farmed populations from wild ones^27^.

In the present study, outlier SVs significantly affected the protein-encoding sequence of at least three genes involved in adaptation to temperature or salinity. In zebrafish, knockout of SLC22A4 results in increased 8-oxoguanine levels showing its key role in eliminating oxygen radicals^28^. Flavin-associated monooxygenases, like the protein encoded by gene SLC22A4, also have antioxidant activity^29,30^. Guanine deaminase (ENSDLAG00005016346) is a key enzyme in purine metabolism, which has been directly linked to temperature adaptation in fish^31,32^. The gene encoding guanine deaminase is also an example of the complex mechanisms that might underlie SVs effects. A deletion encompassing part of the first exon of one isoform and part of a highly specific alternative promoter likely affects where different transcripts of a gene are expressed. The large deletion involving three open chromatin regions and one promoter, all overlapping an LTR-retrotransposon, confirmed that transposable elements might have an important role in the evolution of gene expression regulation as reported in previous studies^33^, by mobilizing regulatory elements. Remarkably, all four elements affected by the SV showed high tissue-specificity, being active only in mature male gonad, potentially affecting reproduction in a lineage-specific manner. Recent evidence has shown an altered sex ratio, with a predominance of males, in an admixed Western Mediterranean population^21^. It should be noted, however, that all individuals from both lineages analysed in our study were male, so a confounding effect between lineage and sex cannot be excluded. This limitation is likely unavoidable given the sampling design used in Duranton et *al*^14,16^. However, it is highly unlikely that the large number of highly divergent genomic regions identified here are all linked to loci involved in sex determination. In fact, a recent study found a single significant sex-tendency QTL on LG19 (∼9.1 Mb region, containing gsdf) exclusively in the Atlantic lineage-based population, with no comparable signal in either Mediterranean population^34^.

Recombination should be accounted for when analyzing genomic divergence, as linkage disequilibrium will affect any outlier analysis. Caution should be exerted when looking at a single outlier locus and trying to discuss its potential role in genomic divergence. When selection acts on a locus, linked neutral or weakly selected sites nearby can “hitchhike” along, leading to local reductions in gene flow and elevated differentiation at both the target locus and its genomic neighborhood^12^. Therefore, functional interpretations of selection acting on single loci might be misleading. On the other hand, genome and chromosome-wide association-enrichment analysis between local genomic divergence and functional features should be largely unaffected by genetic hitchhiking, unless local recombination rates correlate with the presence of specific functional features. Significant heterogeneity in local recombination rates has been reported across major taxonomic assemblages^35–37^. Variation in local recombination rates has been already reported in the European sea bass^17^, with crossing-over occurring more frequently at the chromosome ends as observed in other teleost species^38^, and further confirmed by the present study (**Additional file 2: Fig. S2**).

It has been recently suggested that functional interactions between regulatory elements within the same chromosome lead to lower recombination across the regions of the human genome where such regulatory elements are located, creating “recombination rate valleys”^39^. On the other hand, another study found no significant concordance between chromatin interactions and linkage disequilibrium, even at short (<25 kb) genomic distances^40^. However, there is no evidence for “recombination rate valleys” linked to functional features in the sea bass genome. There is actually clear indication for chromosome 17 that the presence of both genes and all types of regulatory elements is positively correlated with higher recombination rates (**Additional file 1: Table S7**). Such positive correlations have been reported in dogs, snakes, birds, and other fish species^41–43^, attributed to more frequent crossing-overs occurring at accessible chromatin regions, especially promoters. The positive association between functional features and recombination rate at chromosome 17 certainly deserves further attention in future work. However, with the exception of chromosome 17, there was no correlation between the density of regulatory features or genes and local recombination rates. This suggests that the highly significant association between either outlier SNPs or local high Fst values and the density of promoters and enhancers cannot be explained by recombination rate variation. As mentioned, hitchhiking might lead to overestimating the number of loci directly under divergent selection, although relative enrichment tests and correlation analyses should not be significantly affected, unless the presence of genomic features being correlated with divergence is also correlated with local recombination rates.

The significant enrichment of outlier SNPs in gene bodies and regulatory elements hints that genomic divergence between Atlantic and Mediterranean lineages cannot be explained solely by neutral evolution^12^. Prior to any tentative explanations, it should however be noted that outlier SNPs are only potentially enriched in regulatory regions, with no guarantee that these regions are actually universally active. This is because the individuals in the reference genome, and thus the epigenomic map, differed from the ones employed in the analysis of natural populations. The two most likely hypotheses are either that the past separation between the two marine basins during the last glaciation led to the establishment of BDM incompatibilities or that diverging environmental conditions such as water temperature, salinity, and pathogens, especially at the extremes of the sea bass distribution range, promoted adaptation to local conditions through positive selection. These two hypotheses are not mutually exclusive and it is generally difficult to assess which type of selection is the most relevant. Both scenarios can explain the enrichment of outlier SNPs in regulatory elements. On the other hand, it can be reasoned that BDM incompatibilities should not show a particular bias toward regulatory elements that are active either in early development or in adult tissues. The significant enrichment of outlier SNPs in enhancers active just in adult tissues might be better explained by selection on regulatory variants active in adult fish. This interpretation needs caution, as observations in the adult epigenome reference tissue could not capture the entire range of possible regulatory activity in natural populations. Nonetheless, it is worth noting that regulatory elements active in adult stages are more frequently the target of positive selection compared to promoters and enhancers involved in development, as the latter experience strong purifying selection^44^. The role of regulatory divergence in local adaptation, as opposed to its role in BDM incompatibilities, remains largely unexplored^10^. Our findings suggest regulatory elements as a driver of local adaptation, as already observed in other teleosts, with cis-regulatory variants driving ecological divergence in Lake Masoko cichlids^45^, a system in which genomic islands of differentiation enriched in adaptive genes were found^46^. A specific type of genomic incompatibility, mitonuclear incompatibility, has been recently shown to have an important role in reducing hybrid fitness and generating species boundaries^47,48^. It is associated with mismatched mitochondrial protein combinations encoded respectively by the nuclear genome and the mitochondrial one in hybrids. Enrichment analysis of outliers in nuclear genes encoding mitochondrial proteins showed no evidence in support of mitonuclear incompatibilities between Atlantic and Mediterranean sea bass.

Another possible line of evidence to discriminate between BDM incompatibilities and selection is to test whether outlier loci in regulatory elements show a significant functional enrichment in biological processes and molecular functions linked to abiotic and biotic factors characterizing the two marine basins. The underlying rationale is that BDM incompatibilities are not expected to be directly linked to adaptation to local conditions. However, it is quite challenging to associate regulatory elements to specific genes and their functions, since enhancers often regulate the expression of genes located at variable distances and promoters might also act as enhancers^49^. On the other hand, most interactions between regulatory elements and their target genes occur within chromosomes, therefore chromo-some-level functional enrichment analysis should capture most of these interactions. Functional enrichment analysis of genes located in the three chromosomes displaying strong positive correlation between local Fst values and regulatory element coverage (chromosomes 1, 9, 18, **Additional file 1: Table S3**) returned highly enriched molecular functions and biological processes (**Fig. 4**) that represent promising candidates for local adaptation in the European sea bass. Amino acid – sodium/chloride ions symporters (chromosome 1) are considered highly relevant in salinity adaptation^30,50^. Positive regulation of immune responses (chromosome 9) aligns with the demonstrated gradient in resistance to viral disease across geographic sea bass lineages^19,20^. Flavin-containing monooxygenases are important players in the response to oxidative stress linked to both thermal and salinity varia-tion^29,30^. The use of a chromosome-level enrichment approach is also supported by the known clustering in the same genomic regions at the chromosome level of genes involved in the similar biological process or encoding proteins with related molecular functions. The location of genes across the genome are not completely random, with functionally related genes frequently located on the same chromosome in close spatial proximity, which may act to coordinate their transcriptional regulation. Evidence from the analysis of outlier SVs affecting protein function and gene expression appeared to be quite concordant with observations for SNPs. In fact, lineage-specific deletions targeted genes that encode for proteins involved in oxidative stress response, flavin-associated monooxygenase and organic cation transporter, or in temperature adaptation.

## Conclusions

Extensive functional annotation of the European sea bass genome provided a novel epigenome landscape to overlay with signatures of population genomic divergence. This allowed us to observe highly significant correlations between local population divergence and regulatory elements, in particular enhancers. Overall, there is clear evidence for a major role of genetic variation at these regulatory regions in the evolution of divergent lineages across the Atlantic-Mediterranean divide. Such evidence is robust against possible disturbing factors like variation in local recombination rates and divergence hitchhiking. Detailed analysis of the distribution and functional enrichment of regulatory elements associated with genomic divergence suggests that positive selection for adaptation to local conditions acting on a large number of loci across the genome might have contributed to deep divergence of sea bass lineages in the absence of strong present-day barriers to gene flow.

## Methods

### Genome sequencing and assembly

A containerized pipeline for the de-novo genome was used to assemble the European sea bass (*Dicentrarchus labrax*) genome. The input was Oxford Nanopore Technologies (ONT) long reads and Illumina short reads. The Illumina data was sequenced from populations of three fish farms in Greece. These farmed populations were expected to have reduced heterozygosity compared to wild populations. The assembly pipeline employed a series of software to carry out the assembly and polishing. First, Trimmomatic v0.32^51^ was used for read trimming, while Nanoplot v1.47.0^52^ and FastQC v0.12^53^ were used for ONT and Illumina sequence data quality control, respectively. Subsequently, the Flye assembler v2.9.6^54^ built an initial assembly using the ONT data, while Racon v1.5.0^55^ and Medaka v2.2.2 (https://github.com/nanoporwww.bioinformatics.babra-ham.ac.uk/projects/fastqcetech/medaka) were used for an initial polishing round using the ONT data. Next, Pilon v1.24^56^ was used with the short-read data for a final polishing step. Quast v5.3.0^57^ and BUSCO v.6^58^ were used assess the quality of produced assemblies in terms of contiguity and completeness respectively. The final assembly obtained was scaffolded to a near chromosome level, guided by the published assembly of Tine et *al*.^15^, using RagTag v2.1.0^59^ to map and scaffold our assembly contigs based on the published scaffolds.

### Epigenome annotation: embryonic and adult tissue samples

A batch of fertilized European sea bass eggs was provided by Valle Cà Zuliani Società Agricola srl (Pila di Porto Tolle, Rovigo, Italy) immediately after fertilization. Eggs were housed in an aerated aquarium with saltwater at the facilities of the Department Comparative Biomedicine and Food Science of the University of Padova at 15.0°C (±0.5°C) until hatching. A total of 6 developmental stages were collected for ATAC-seq and ChIP-seq library preparation (3 biological replicates per stage), namely i) Early gastrula, ii) Mid-gastrula (60% epiboly), iii) Early (10%) somitogenesis, iv) Mid (50%) somitogenesis, v) Late (90%) somitogenesis and, vi) Near-to-hatching stage. All these procedures were part of a large annotation effort under the framework of the EU project AQUA-FAANG (https://www.aqua-faang.eu/). Detailed protocols for sample collection and egg dechorionation are available in the FAANG repository (data.faang.org; https://data.faang.org/api/fire_api/experiments/UNIPD_SOP_Embryos Dechorionation and Chorion Constituents Cleaning Dlab_20201012.pdf).

A total of 12 European sea bass obtained from a commercial broodstock (Valle Cà Zuliani Società Agricola srl, Pila di Porto Tolle, Rovigo, Italy) were transferred to the Istituto Zooprofilattico Sperimentale delle Venezie (IZSVe, Legnaro, Padova, Italy). The experimental group comprised 6 sexually immature individuals (approximately 22 months old, total length 29.8±1.7 cm, weight 261.9±44.2 gr) and 6 sexually mature individuals (approximately 42 months old, total length 36.7±3.1 cm, weight 488.3±119.9 gr). For each reproductive stage, the group consisted of 3 males (‘IM’ = immature male, ‘MM’ = mature male) and 3 females (‘IF’ = immature female, ‘MF’ = mature female). Fish were maintained in indoor tanks filled with artificial saltwater (30‰ salinity, temperature 19 ± 1°C, oxygen 6 ppm) and exposed to an artificial photoperiod (14 h of light, 10 h of darkness). Before tissue collection, fish were anesthetized by bath (MS-222; 100mg/L) and consequently euthanized with anaesthetic overdose (MS-222; 150 mg/L). Five different tissues (skeletal muscle, gill, brain, liver and gonads) were collected per animal, immediately frozen in dry ice and stored at −80 °C until ATAC-seq and ChIP-seq library preparation.

All animal procedures were conducted in compliance with Directive 2010/63/EU of the European Parliament and of the Council and were implemented in accordance with the Italian Legislative Decree of 4 March 2014, No. 26, authorised by the Italian Ministry of Health (authorisation no. 641/2018-PR).

### Epigenome annotation: ATAC-seq library preparation and sequencing

ATAC-seq libraries were prepared from IM, MM, IF and MF brain, liver, gill, skeletal muscle, and gonad samples (n=3 libraries per tissue, gender and reproductive stage) using Omni-ATAC^60^ with modifications (details here: https://data.faang.org/api/fire_api/experiments/NMBU_SOP_OmniATAC_protocol_20200429.pdf). Briefly, frozen tissue fragments were homogenized using a pre-chilled 2-ml dounce tissue homogenizer containing homogenization buffer. Connective tissue and residual debris were removed using a a 70-μm nylon filter. Intact nuclei were collected by density gradient centrifugation over iodixanol (25% iodixanol, 29% iodixanol, 35% iodixanol). After PBS washing, nuclei were counted by trypan blue staining and 50,000 nuclei were used in the transposition reaction with Tn5 transposase using the Illumina Tagment DNA Enzyme and Buffer kit (Illumina, San Diego, California, USA). Transposition reactions were cleaned up with MinElute PCR Purification Kits (Qiagen, Hilden, Germany) and uniquely barcoded libraries were obtained following amplification using Illumina Nextera DNA Unique Dual Indexes (Illumina).

ATAC-seq libraries were also prepared for the 6 developmental stages described above (n=3 each; total 18 libraries in total) (details here: https://data.faang.org/api/fire_api/experiments/UNIPD_SOP_ATACseq_from_freshly_collected_embryos_20201020.pdf). Briefly, freshly-collected dechorionated embryos (∼50 embryos per biological replicate) were washed in PBS 1× adjusted to sea bass osmolarity (350 mOsm/kg) and mechanically dissociated using a pre-chilled 2-ml Dounce tissue homogenizer containing RSB buffer (350 mOsm/kg). Cell suspensions were filtered through a 70 µm cell strainer, centrifuged, and washed with PBS 1X (350 mOsm/kg). Cells nuclei were counted by trypan blue staining and, for each sample, 50,000 cells were treated with DNase I to reduce genomic DNA contamination prior to nuclei isolation. Nuclei were released by detergent-based lysis using ATAC-RSB-L buffer, washed with ATAC-RSB-W buffer, and collected by centrifugation. Purified nuclei were then employed in transposition reactions were before library preparation as described above for tissues. Final libraries were purified using AMPure XP beads and quality-checked for concentration and size distribution before sequencing.

After quality assessment, all ATAC-seq libraries were sent to Novogene (Cambridge, UK) for 150bp paired-end sequencing on an Illumina NovaSeq 6000 system platform.

### Epigenome annotation: ChIP-seq libraries preparation and sequencing

European sea bass tissues were processed for ChIP-seq library preparation using a modified µChIP-mentation (Diagenode, Seraing, Belgium) protocol optimised for frozen samples. For skeletal muscle, gill, brain and liver tissues, ChIP-seq libraries were generated from two biological replicates for each sex and developmental stage combination (IM, IF, MM, MF; n = 2 per group). For gonadal tissue, three biological replicates were analysed for each sex and developmental stage combination (n = 3 per group). Briefly, tissue fragments were mechanically dissociated using a pre-chilled 2-ml Dounce tissue homogeniser containing ice-cold 1× PBS (350 mOsm/kg) supplemented with protease inhibitors (PIC tablet, Roche). Homogenates were filtered through a 70-µm cell strainer, and the resulting cell suspensions were cross-linked with 37% formaldehyde for 8 min at room temperature. Cross-linking was quenched with glycine, and cells were washed and lysed in complete tL1 buffer. Chromatin was sheared by sonication using a Covaris M220 instrument (Peak Power 75, Duty Factor 5, Cycles/Burst 200) for 7 min at 7 ± 1 °C. An aliquot of sheared chromatin was decross-linked and purified to assess fragment size and concentration. For each immunoprecipitation, 30 ng of sheared chromatin were used. Chromatin immunoprecipitation, tagmentation, library amplification and clean-up were subsequently performed following the µChIPmentation for Histones kit protocol (Diagenode). Four histone modifications were investigated: H3K4me3 (promoter-associated regions), H3K27ac (active enhancer and promoter regions), H3K27me3 (associated with Polycomb repressive complexes), and H3K4me1 (enhancer-associated regions).

For the developmental stages, an average of 50 dechorionated embryos per biological replicate were cross-linked with 37% formaldehyde for 8 min at room temperature. Cross-linking was quenched with glycine, and embryos were washed with 1× PBS and stored at −80 °C until further processing. ChIP-seq libraries were then prepared following the µChIPmentation for Histones protocol (Diagen-ode). Three histone modifications were investigated in developmental stages: H3K4me3 (promoter-associated regions), H3K27ac (active enhancer and promoter regions) and H3K27me3 (associated with Polycomb repressive complexes).

For both the tissue and developmental stage samples, final libraries were purified using AMPure XP beads and quality-checked for concentration and size distribution before sequencing. After quality assessment, all libraries were sequenced by Novogene using the Illumina NovaSeq 6000 platform (paired-end 150 bp), generating approximately 45 million reads per sample/histone mark.

### Epigenome annotation: European sea bass regulatory features

Ensembl’s regulatory annotation was generated by processing paired-end ATAC-seq and ChIP-seq data for H3K4me1, H3K4me3 and H3K27ac to produce functional annotations of promoters, enhancers, and unclassified open chromatin regions^22^. Adapter sequences were removed using NG-merge^61^ with parameters “-a-u 41”. Genomic alignments were performed with Bowtie 2^62^, and peaks were called using Genrich v0.6.1 (https://github.com/jsh58/Genrich). Open chromatin peaks were merged across samples to generate a unified set of genomic features. Features located near the 5’ end of annotated genes were classified as promoters, those overlapping H3K4me1 or H3K27ac peaks were annotated as enhancers, and the rest were retained as unclassified open chromatin regions. Activity of regulatory features by tissue and condition was defined by overlap with sample open chromatin peaks. Further details are available at https://regulation.ensembl.org and in Ilsley et *al*^22^.

### Population genomics: SNP calling

The dataset included 65 wild European sea bass from three populations: 13 males from the Atlantic Ocean (English Channel), 30 individuals from the western Mediterranean Sea (WestMed - Gulf of Lion, consisting of 21 females, 8 males and 1 undefined), and 22 males from the eastern Mediterranean Sea (EastMed - 11 from Turkey and 11 from Egypt) as described in Duranton et *al*^14^ and Duranton et *al*^16^.

The sequence reads were assessed for sequencing quality using FastQC v0.12^53^ and consequently trimmed of any sequencing adaptors and low-quality sequences using Trimmomatic v0.32^51^; high quality reads of length ranging from 70bp to 150bp were retained for variant calling. Burrows-Wheeler Aligner (BWA) v0.7.17^63^ was then used to align the clean reads to the European sea bass genome dlabrax2021 (GCA_905237075.1) downloaded from Ensembl public database v.115. Samtools v1.15 and BCFtools v1.13^64^ were subsequently used for genotype calling. The output file was then filtered with BCFtools to delete indels and non-biallelic SNPs. Regulatory features (i.e., enhancer, promoter and open chromatin regions) of the European sea bass genome dlabrax2021 (GCA_905237075.1) were retrieved from Ensembl public database v.115.

### Population genomics: Structural Variant (SV) calling

SV calling was performed using a pipeline that has been used to produce high-confidence SV landscapes in the genomes of farmed teleosts including Atlantic salmon^65^, rainbow trout^66^ and most recently European sea bass^27^, where the SV dataset produced in this study was presented, validated and used for a distinct analysis comparing farmed and wild populations. Farmed individuals were excluded from the present 65-sample wild-population dataset, and SV genotyping quality was re-assessed specifically for the Atlantic/EastMed contrast. Briefly, SV detection and genotyping were performed using Lumpy v0.2.7^67^ and SVtyper v0.7.0^68^ within the Smoove framework (https://github.com/brentp/smoove). SV filtering was conducted following the procedures described in Jiao et *al*^27^. Specifically, SVs were removed if they: i) were detected on unplaced scaffolds; ii) represented translocations; iii) overlapped scaffold gaps or were located in genomic regions with sequencing depth >100×); iv) were classified as low-quality SVs by Duphold v0.2.1^69^; or (v) had an allele count ≤2 and/or a genotype call rate <80%. All SVs passing these filters were further subjected to visual inspection using SV-plaudit^70^. Only SVs retained after SV-plaudit validation were considered high-confidence SVs (n=38,408) and included in the final dataset, consistent with previous studies^27,65^. This high-confidence SV dataset was also shown to recover the expected population genetic structure of the three wild sea bass populations^27^.

### Population genomics: Sea bass linkage map

The first step in determining recombination rates was the construction of linkage maps using Lep-MAP3^71^. Pedigree and genotype data from a full-factorial mating design (25 dams x 25 sires; 990 offspring), comprising 29,888 SNPs generated via the MedFish SNP array^72^, were retrieved from Mukiibi et *al*^20^. A total of 447 full-sib families were employed, each consisting of 1 to 6 offspring per family. The module *ParentalCall2* was set with option halfSibs = 1, to allow inference of half-sib parental genotypes. In module *Filtering2*, the default p-value limit of 0.001 for segregation distortion was kept (dataTolerance = 0.001), retaining 27,930 SNP. The module *SeparateChromosomes2* then assigned them into 24 main linkage groups (LGs), joining markers based on a pairwise LOD score higher than 68 (lodlimit = 68). Marker ordering and centiMorgan (cM) position estimations were performed with the module *OrderMarkers2* using default settings (useMorgan = 1). Final linkage maps were plotted with the R package *LinkageMapView*^73^.

To estimate the local recombination rate, we compared our newly constructed genetic map to the physical positions from the European sea bass reference genome (Ensembl release 115) using MAREYMAP^74^. The LOESS method was used to determine the local recombination rate in cM per Mb. The median recombination rate per 0.1-Mb sliding windows has then been estimated for comparison with the median of the fixation index (Fst) distribution (see below).

### Population genomics: genetic structure analysis

Genetic structure was examined with fastSTRUCTURE v1.0^23^, which infer the ancestral individual proportions of genetic membership to the different populations^23^. We evaluated the model for clusters (k) ranging from 1 to 5 to determine the optimal number of populations that best represent the underlying genetic structure. The prior parameter was set to logistic (--prior = logistic), allowing for population-specific allele frequencies modeled using a normal logistic distribution. This option, although computationally demanding, is more accurate compared to the default simple prior, and suited for small sample sizes (42 Raj et *al*., 2014). Results were visualized using the *pophelper* package in R v4.4.1^75^.

To investigate genomic differentiation, the West Mediterranean population was excluded from the analysis enabling a direct comparison between the theoreticaly more divergent Atlantic and East Mediterranean individuals. This exclusion was necessary as the West Mediterranean group exhibited significant introgression from the Atlantic population, which could hinder the accurate identification of genomic islands. Genomic regions under selection were identified by comparing the Atlantic and East Mediterranean populations through the *pcadapt* R package^76^. Two principal components (K = 2) were considered representative of the genetic structure, and a minor allele frequency threshold of 0.10 (min.maf = 0.10) was applied for p-value calculation. Markers with p-values below 1% α (α = 0.01) threshold were identified as outliers, following a Bonferroni correction with the R package *stats*.

The fixation index (Fst), estimated using VCFtools^77^, was employed to further characterize the degree of genetic differentiation between the Atlantic and East Mediterranean populations. The distribution of genomic islands across chromosomes was computed as the median Fst value within non-overlapping sliding windows of 100 Kb.

### Population genomics: statistical analyses

A functional enrichment analysis was conducted on the outliers (both SNPs and SVs) identified in the genomic divergence between the Atlantic and East Mediterranean populations. Specifically, we investigated the over-or under-representation of these variants across different regulatory elements and developmental stages. These categories included intergenic variants, regulatory variants (further subdivided into open chromatin, enhancers, and promoters), downstream/upstream variants, transcript variants (separated into intron variants, splice region variants, 5’ UTR/3’ UTR variants), tissue-specific variants (active in mature or immature tissues), pleiotropic variants (active in multiple tissues), developmental stage variants, synonymous and nonsynonymous variants. For each category, the number of outliers variants was compared with the variants from the remainder of the dataset by constructing contingency tables. Statistical significance of association was evaluated using Pearson’s chi-square test or Fisher’s exact test, all performed in the R package *stats*.

To detect whether specific functional regions contribute to patterns of genomic differentiations, several metrics were quantified and compared across 100-kb windows. For the whole genome and for each chromosome, we calculated gene coverage, regulatory feature coverage (enhancers, promoters, and open chromatin regions), SNP and SV densities, and median recombination rates. Pairwise associations among these metrics and median Fst values were evaluated using Spearman correlation tests, implemented in R with the package *stats*.

Gene set enrichment analyses were performed using Gene Ontology (GO) terms and KEGG pathways through g:Profiler v.e111_eg58_p18_f463989d, applying the g:SCS multiple testing correction method with a significance threshold of 0.05^78^. First, genes associated with all SNP outliers were tested for enrichment of gene sets relative to genes associated with background SNPs. Subsequently, gene set enrichment analyses were conducted for genes located exclusively on chromosomes 1, 9, and 18, each compared against all genes located in the rest of the genome. These chromosomes were identified as potentially relevant based on the functional enrichment analyses described above.

To test whether mitonuclear incompatibilities could influence genomic divergence between sea bass populations, a specific enrichment analysis was performed. Using the *MitoCarta3.0* database^79^ and the g:Orth function of g:Profiler v.e111_eg58_p18_f463989d, orthologous genes between humans and sea bass were identified. We then evaluated the potential overrepresentation or underrepresentation of nuclear genes encoding mitochondrial proteins among SNP outliers relative to SNP background.

## Supplementary Information

- File name: Additional file 1

Format:.xlsx

Title: Table S1

Description: Mean and median percentage of chromosome regions annotated as promoters, enhancers, and open chromatin

- File name: Additional file 1

Format:.xlsx

Title: Table S2

Description: Contingency table showing outlier and background genes with respect to *Mito-Carta3.0*

- File name: Additional file 1

Format:.xlsx

Title: Table S3

Description: Spearman correlation between coverage of regulatory features and median Fst in 100kb windows

- File name: Additional file 1

Format.xlsx

Title: Table S4

Description: Spearman correlation between SNP density and functional features coverage

- File name: Additional file 1

Format.xlsx

Title: Table S5

Description: Spearman correlation between SV and functional features coverage

- File name: Additional file 1

Format.xlsx

Title: Table S6

Description: Overall and chromosome-by-chromosome correlation analysis between local Fst and recombination rate

- File name: Additional file 1

Format.xlsx

Title: Table S7

Description: Spearman correlation analysis of coverage by functional features and local recombination rates

- File name: Additional file 2

Format:.pdf

Title: Fig. S1

Description: Results of Structure analysis

- File name: Additional file 2

Format:.pdf

Title: Fig. S2

Description: Median recombination rate estimates (in cM/Mb) along the 24 chromosomes of the European sea bass

## Acknowledgements

We are grateful to Gil Rosenthal, Flavia Termignoni-Garcia, and François Allal for reviewing a final version of the paper.

## Authors’ contributions

L.B. and D.J.M. designed and supervised the project. V.P., T.M., J.K., and C.S.T. conducted the genome sequencing and assembly. M.B. and D.B. performed embryonic and tissue samples collection for the epigenome annotation. S.F. and R.F. prepared ATAC-seq and ChIP-seq libraries and sequencing. G.R.I. provided the functional build for european sea bass regulatory features. M.B. conducted the SNP calling. Z.J. performed the SV calling. A.L. and M.B. performed the linkage map, genetic structure and statistical analyses. A.L., S.F. and L.B. contributed to the writing manuscript with inputs from all the other authors. All authors read and approved the final manuscript.

## Availability of data and materials

ATAC-seq and ChIP-seq sequence data generated in this study have been deposited in the EMBL-EBI repository under the accession numbers PRJEB52775 (ATAC-seq for European Sea bass tissues), PRJEB59432 (ChiP-seq for European Sea bass tissues), PRJEB52159 (ATAC-seq for European Sea bass embryos), and PRJEB59434 (ChIP-seq for European Sea bass embryos).

## Declarations

### Ethics approval and consent to participate

Not applicable.

### Consent for publication

Not applicable.

