## Additional file 2 for "The epigenomic landscape of deep lineage divergence: The case of the European sea bass"

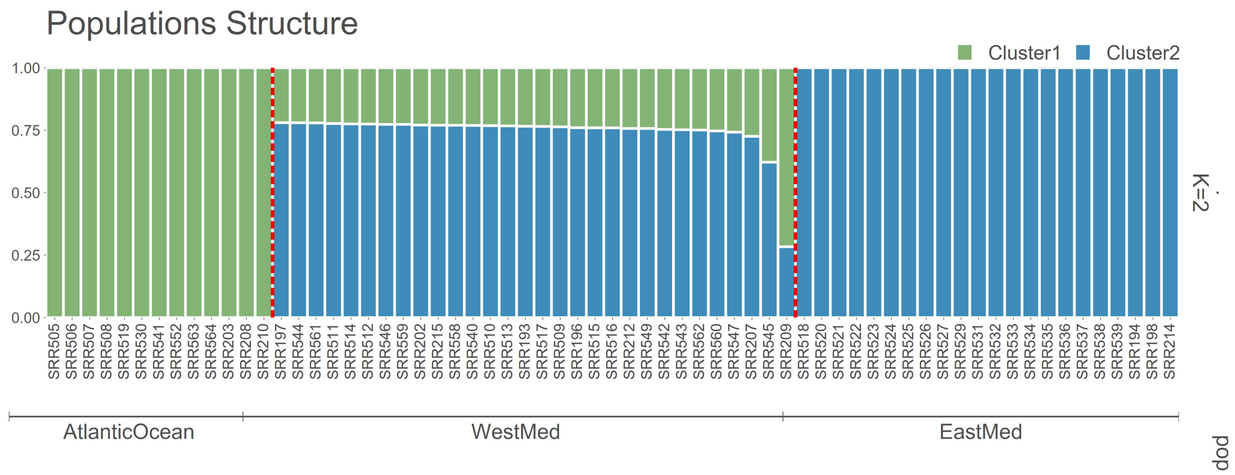


**Fig. S1 Results of Structure analysis**

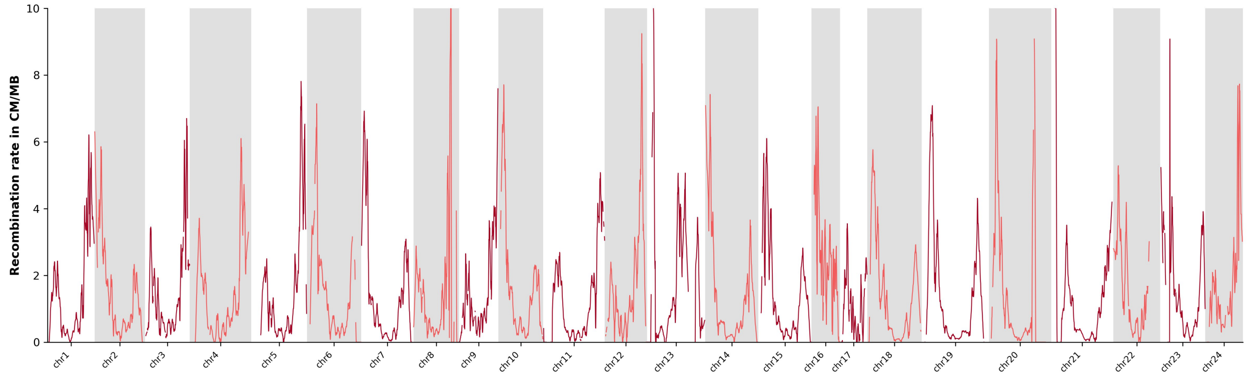


**Fig. S2 Median recombination rate estimates (in cM/Mb) along the 24 chromosomes of the European sea bass.**
